## Supplementary material for "Introducing the CRISPR/Cas9 cytosine base editor toolbox ‘LeishBASEedit’ – Gene editing and high-throughput screening in *Leishmania* without requiring DNA double-strand breaks, homologous recombination or donor DNA": Protocol S1.pdf

Julius-Maximilians-
UNIVERSITÄT
WÜRZBURG
:::::: Biozentrum
Zell- und Entwicklungsbiologie
.ZOO

www.biozentrum.uni-wuerzburg.de/zeb/team/staff-scientists/tom-beneke/

LeishBASEedit

**Dr. Tom Beneke**

Cell and Developmental Biology
Biocenter, University of Wuerzburg
Am Hubland • D-97074 Würzburg


### Cloning guide target sequences into base editor plasmids for *Leishmania* editing
#### Protocol version October 2022

#### Workflow

Primer pairs given by LeishBASEedit can be used to clone guides into BbsI sites of base editor plasmids, such as pLdCH-hyBE4max. The general workflow is shown below. First, both oligos (primer sequences given by LeishBASEedit primer design) are annealed and phosphorylated, before they are being ligated into a dephosphorylated vector that was digested with BbsI. The resulting plasmid can be transfected into *Leishmania* species.

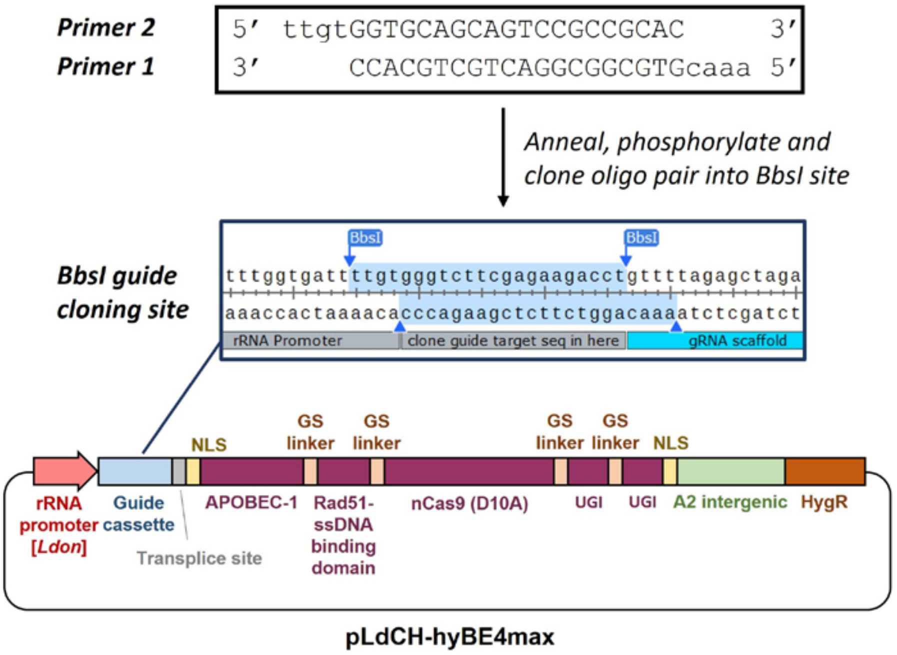

#### Step-by-step protocol

1. Digest and Dephosphorylate base editor plasmid
   - 5 µg plasmid
   - 2 µl BpiI (BbsI) [ThermoFisher, ER1011]
   - 2 µl FastAP [ThermoFisher, EF0651]
   - 3 µl 10x Tango Buffer [ThermoFisher]
   - 30 µl total volume with dd$H_2O$
   - Incubate at 37°C for at least 4 hours or better overnight
   - PCR purify digested and dephosphorylated plasmid (confirm linearization of plasmid on gel)

2. Anneal and phosphorylate oligo pair
   - Order standard desalted oligos (25 nmole scale)
   - 1 µl oligo 1 (100 µM)
   - 1 µl oligo 2 (100 µM)
   - 1 µl T4 DNA Ligation Buffer (ATP needed for phosphorylation)
   - 0.5 µl T4-Polynukleotid-Kinase [ThermoFisher, EK0031]
   - 6.5 µl dd$H_2O$
   - Phosphorylate at 37°C for 30 minutes, followed by 5 minutes at 95°C
   - Then anneal by ramping down from 95°C to 25°C at 5°C/minute
   - Dilute (1:200) annealed and phosphorylated oligo pair by mixing 1µl with 199µl dd$H_2O$

3. Ligation and transformation
   - 50ng digested and dephosphorylated plasmid (from Step 1)
   - 1µl diluted oligo pair (from Step 2)
   - 1µl T4 DNA Ligation Buffer
   - 0.5µl T4 DNA Ligase [ThermoFisher, EL0014]
   - 10µl total volume with dd$H_2O$
   - Incubate at 37°C for 2 hours
   - Transform entire volume into 50-100 µl competent cells (e.g. TOP10, Stbl3, XL1Blue)
   - Spread out on AmpR plate and compare to “no-insert” control
