## Supplementary material for "Introducing the CRISPR/Cas9 cytosine base editor toolbox ‘LeishBASEedit’ – Gene editing and high-throughput screening in *Leishmania* without requiring DNA double-strand breaks, homologous recombination or donor DNA": Protocol S2.pdf

Julius-Maximilians-UNIVERSITÄT WÜRZBURG

Biozentrum
Zell- und Entwicklungsbiologie
.ZOO

LeishBASEedit

www.biozentrum.uni-wuerzburg.de/zeb/team/staff-scientists/tom-beneke/

**Dr. Tom Beneke**

Cell and Developmental Biology
Biocenter, University of Wuerzburg
Am Hubland • D-97074 Würzburg


### Transfecting base editor guide plasmids into *Leishmania*
### Protocol version October 2022

#### Step-by-step protocol

Base editor plasmids that contain guide target sequences can be transfected into *Leishmania* species following conditions outlined in Schumann Burkard et al., Mol Biochem Parasitol 2011; 175, 91-94.

<u>Transfection Buffer</u>

1. Prepare the following buffers:

**<u>3x Tb-BSF buffer stock</u>**
200 mM $Na_2HPO_4$
70 mM $NaH_2PO_4$
15 mM KCl
150 mM HEPES pH 7.4

**<u>$CaCl_2$ stock</u>**
1.5 mM $CaCl_2$ in water

<u>Cell-Plasmid-Transfection-Mix</u>

1. Dilute 5-10 µg of plasmid DNA into 50 µl $ddH_2O$ and heat sterilize for 5 minutes at 95°C
2. For each transfection (see transfection protocol below) a Cell-Transfection-Mix is prepared by mixing:
    - 25 µl $CaCl_2$
    - 83 µl 3x Tb-BSF
    - 92 µl $ddH_2O$ and

<u>Transfection protocol</u>

1. Prepare an exponentially growing culture of *Leishmania*
2. Prepare the required volume of transfection buffer according to instructions above (ensure all solutions are sterile)
3. Pre-warm medium and collect the required number of cells (at least 5E6 cells per transfection) at 800g for 5 minutes
4. Take off supernatant and wash in 1 – 5 ml of Cell-Transfection-Mix (as specified above)
5. Resuspend pellet in desired final volume of Cell-Transfection-Mix
6. Mix 200 µl Cell-Transfection-Mix with 50 µl heat sterilized plasmid DNA and transfer into electroporation cuvette. (Note: Leave the cells in the cuvettes for the shortest possible time. When doing a big batch of transfections, do no more than five cuvettes at a time and keep the remaining cells in the Eppendorf tube until use)
7. Transfection: one pulse with X-001 (Amaxa Nucleofector 2b)
8. Quickly and carefully transfer cells into 5 ml pre-warmed medium (cell concentration should not be higher than 1E6 cells/ml) and rinse cuvette once with medium.
9. Leave cells to incubate for 8-16 h, then add the required selection drug and incubate until drug resistant populations emerge
