## Supplementary material for "Introducing the CRISPR/Cas9 cytosine base editor toolbox ‘LeishBASEedit’ – Gene editing and high-throughput screening in *Leishmania* without requiring DNA double-strand breaks, homologous recombination or donor DNA": LeishBASEedit_v1.sh

```
#!/bin/bash
#-------------------------------------------------------------------------------------------------------------------------------------------------------------------------------------------------------------------------
#README:
#This script was written by Tom Beneke and designs guides to introduce STOP codons within the first 50% of any given ORF.
#These CBE guides are designed using an autocbei pipeline (CRISPR-CBEI, Yu et al., mSystems 2020) 

#Dependencies:
#This script uses autocbei (https://github.com/atlasbioinfo/CRISPR-CBEI/tree/master/autocbei)
#install autocbei with: $ pip install autocbei

#Instructions on usage:
#Place this script "LeishBASEedit_v1.sh" in a single folder together with:
#(1) your reference genome file (chromosomes or contigs, in fasta format).
#(2) your CDS file (in fasta format).
#(3) File: "hyBE4max_specifics.csv" (determines the editing window parameters for the used CBE and can modified if desired).
#Navigate to your directory and execute the script by "sh LeishBASEedit_v1.sh"

#Output:
#The main output file is named: $REFERENCE_GENOME"_LeishBASEv1_output.txt
#Details of each column in this output file can be found on LeishBASEedit.net
#In addition, there are several processing and raw data files that may be needed for troubleshooting.

#Testing the script on your system
#To test the script we have provided two test files:
#(1) An CDS file ("script-test-CDS-file.fasta")
#(2) A genome reference test file ("script-test-genome-reference-file.fasta")
#Testing the script with these two files will take approximately 5 minutes.

#Program start
#-------------------------------------------------------------------------------------------------------------------------------------------------------------------------------------------------------------------------
pause(){
   read -p "Press [Enter] key to continue..." fackEnterKey
}

pauseRUN(){
   read -p "Press [Enter] key to start primer design with LeishBASEedit v1.0..." fackEnterKey
}

echo "Welccome to LeishBASEedit primer design (version 1.0)"
read -p 'Please enter your reference genome (in fasta format): ' REFERENCE_GENOME
	echo "You are uploading <"$REFERENCE_GENOME"> as a reference genome."
	pause

read -p 'Please enter your CDS file (in fasta format): ' CDS_input
	echo "You are uploading <"$CDS_input"> as a CDS file."
	pauseRUN

#Inputs
echo "We are processing <"$REFERENCE_GENOME"> as a reference genome."
echo "We are processing <"$CDS_input"> as CDS file."

echo "Processing started..."


#CBEI search
echo "Started autocbei guide search"
autocbei -bef hyBE4max_specifics.csv $CDS_input
cd CBEIRaw
cp * ../
cd ..

rm -r CBEIStat
rm -r CBEIPlot

mkdir Processing_files

#filter within first 50% and hihglight number of edits per guide
echo "Extraction of guides within first 50% of CDS"
awk '$3<0.5' hyBE4max4-12_-bef.cbei > Processing_files/hyBE4max4-12_-bef.0.5.txt
awk '{print $7}' Processing_files/hyBE4max4-12_-bef.0.5.txt | awk -F "," ' { print NF-1 } ' > Processing_files/hyBE4max4-12_-bef.stops.txt
echo "Highlighting guides with more than one edit"
paste Processing_files/hyBE4max4-12_-bef.0.5.txt Processing_files/hyBE4max4-12_-bef.stops.txt | awk -F"\t" '{print $1"_"$6"\t"$1"\t"$2"\t"$3"\t"$4"\t"$5"\t"$6"\t"$7"\t"$8"\t"$9"\t"$10"\t"$11"\t"$12}' > Processing_files/hyBE4max4-12_-bef.0.5.stops.txt

#highlight guides with 4-8, 4-10 and 4-12 window and prepare guide score
echo "Highlighting guides with editing window 4-8"
awk -F"\t" '{print $1"_"$6}' hyBE4max4-8_-bef.cbei > Processing_files/hyBE4max4-8_-bef.txt
awk 'NR == FNR{a[$1] = $0;next}; {print $0, $1 in a?a[$1]: "0"}' OFS='\t' Processing_files/hyBE4max4-8_-bef.txt Processing_files/hyBE4max4-12_-bef.0.5.stops.txt | awk 'BEGIN { OFS = "\t" } {if ($14 ~ /./) ($14 = 1)} 1' > Processing_files/hyBE4max4-12_-bef.4-8.txt

echo "Highlighting guides with editing window 4-10"
awk -F"\t" '{print $1"_"$6}' hyBE4max4-10_-bef.cbei > Processing_files/hyBE4max4-10_-bef.txt
awk 'NR == FNR{a[$1] = $0;next}; {print $0, $1 in a?a[$1]: "0"}' OFS='\t' Processing_files/hyBE4max4-10_-bef.txt Processing_files/hyBE4max4-12_-bef.4-8.txt | awk 'BEGIN { OFS = "\t" } {if ($15 ~ /./) ($15 = 1)} 1' > Processing_files/hyBE4max4-12_-bef.4-8.4-10.txt

rm -r *.cbei

#sorting by guide score
echo "Finalizing score"
awk 'BEGIN { OFS = "\t" } {if ($3 == "Minus") ($1 = $2"_0\t"$1); else ($1 = $2"_1\t"$1)} {if ($4 <= 0.2) ($15 = $15"\t1"); else ($15 = $15"\t0")} {if ($4 <= 0.4) ($15 = $15"\t1"); else ($15 = $15"\t0")} 1' Processing_files/hyBE4max4-12_-bef.4-8.4-10.txt > Processing_files/hyBE4max4-12_-bef.4-8.4-10.minpl.txt
awk -F"\t" '{print $1"_"99-($14+$15+$16+$17+$18)"_"$5"_"$8"\t"$3"\t"$4"\t"$5"\t"$6"\t"$7"\t"$8"\t"$9"\t"$10"\t"$11"\t"$12"\t"$13"\t"$14"\t"$15"\t"$16"\t"$17"\t"$18"\t"($14+$15+$16+$17+$18)}' Processing_files/hyBE4max4-12_-bef.4-8.4-10.minpl.txt > Processing_files/hyBE4max4-12_-bef.4-8.4-10.score.txt
sort -k 1 Processing_files/hyBE4max4-12_-bef.4-8.4-10.score.txt | awk 'BEGIN{print "GuideID\tGeneID\tStrand\tPosition\tGuideTarget\tGuideSeq\tGuideCoordinates\tStopCoordinates\tEditingWindow\tPAMseq\tPAMposition\tEditSeq\t#AdditionalStops\tEditWindow_4-8\tEditWindow_4-10\tTargetWithinFirst_20%_of_CDS\tTargetWithinFirst_40%_of_CDS\tTotalScore"}1' > Processing_files/hyBE4max4-12_-bef.4-8.4-10.score.sorted.txt

#Counting Number of guides per gene
echo "Counting Number of Guides per Gene"
awk '{print $2}' Processing_files/hyBE4max4-12_-bef.4-8.4-10.score.sorted.txt | uniq -c > Processing_files/guides-per-gene-count.txt

#Preparing Genome for on and off-target searches
echo "Preparing genome for on-target search"
WORKING_GENOME=working_genome_sgl.fasta
WORKING_GENOME_REVERSE=working_genome_reverse_sgl.fasta

sed -i -e 's/\r$//' $REFERENCE_GENOME

awk '/^>/ {printf("\n%s\n",$0);next; } { printf("%s",$0);}  END {printf("\n");}' $REFERENCE_GENOME | tail -n +2 > Processing_files/$WORKING_GENOME
echo "Making reverse complement of genome"
cat Processing_files/$WORKING_GENOME | while read L; do  echo $L; read L; echo "$L" | rev | tr "ATGC" "TACG" ; done > Processing_files/$WORKING_GENOME_REVERSE


#Check for 0M counts forward
PIPE1="awk '{print "
PIPE2="}'"
PIPE3='"'
echo "Checking 0M counts forward"
mkdir Processing_files/ontarget0M-forward
awk 'BEGIN {print "#!/bin/bash"}' > Processing_files/ontarget0M-forward/M0_genome_count.sh
echo "WORKING_GENOME="$WORKING_GENOME >> Processing_files/ontarget0M-forward/M0_genome_count.sh
cat Processing_files/hyBE4max4-12_-bef.4-8.4-10.score.sorted.txt | awk -v PIPE1="$PIPE1" -v PIPE2="$PIPE2" -v PIPE3="$PIPE3" '{print "grep -o -i "$6$10" Processing_files/$WORKING_GENOME | wc -l | "PIPE1 PIPE3 $1 PIPE3 ", $0" PIPE2" >> Processing_files/ontarget0M-forward/M0_count_f.txt"}' >> Processing_files/ontarget0M-forward/M0_genome_count.sh
sh Processing_files/ontarget0M-forward/M0_genome_count.sh
awk -v OFS="\t" '$1=$1' Processing_files/ontarget0M-forward/M0_count_f.txt > Processing_files/ontarget0M-forward/M0_count_f1.txt

awk 'NR == FNR{a[$1] = $0;next}; {print $0, $1 in a?a[$1]: "@@@@NA@@@@"}' OFS='\t' Processing_files/ontarget0M-forward/M0_count_f1.txt Processing_files/hyBE4max4-12_-bef.4-8.4-10.score.sorted.txt > Processing_files/ontarget0M-forward/pre-final-0MF.txt
echo "Forward 0M count done"

#Check for 0M counts reverse
echo "Checking 0M counts reverse"
mkdir Processing_files/ontarget0M-reverse
awk 'BEGIN {print "#!/bin/bash"}' > Processing_files/ontarget0M-reverse/M0_genome_count.sh
echo "WORKING_GENOME_REVERSE="$WORKING_GENOME_REVERSE >> Processing_files/ontarget0M-reverse/M0_genome_count.sh
cat Processing_files/hyBE4max4-12_-bef.4-8.4-10.score.sorted.txt | awk -v PIPE1="$PIPE1" -v PIPE2="$PIPE2" -v PIPE3="$PIPE3" '{print "grep -o -i "$6$10" Processing_files/$WORKING_GENOME_REVERSE | wc -l | "PIPE1 PIPE3 $1 PIPE3 ", $0" PIPE2" >> Processing_files/ontarget0M-reverse/M0_count_r.txt"}' >> Processing_files/ontarget0M-reverse/M0_genome_count.sh
sh Processing_files/ontarget0M-reverse/M0_genome_count.sh
awk -v OFS="\t" '$1=$1' Processing_files/ontarget0M-reverse/M0_count_r.txt > Processing_files/ontarget0M-reverse/M0_count_r1.txt

echo "Reverse 0M count done"

#joining forward and reverse 0M count
awk 'NR == FNR{a[$1] = $0;next}; {print $0, $1 in a?a[$1]: "@@@@NA@@@@"}' OFS='\t' Processing_files/ontarget0M-reverse/M0_count_r1.txt Processing_files/ontarget0M-forward/pre-final-0MF.txt > Processing_files/pre-final-0M.txt
awk -F"\t" '{print $1"\t"$2"\t"$3"\t"$4"\t"$5"\t"$6"\t"$7"\t"$8"\t"$9"\t"$10"\t"$11"\t"$12"\t"$13"\t"$14"\t"$15"\t"$16"\t"$17"\t"$18"\t"($20+$22)}' Processing_files/pre-final-0M.txt | awk 'BEGIN { OFS = "\t" } {gsub("TotalScore\t0","TotalScore\t0M_count")} 1' > Processing_files/pre-final2-0M.txt

#generating CBE primers
cat Processing_files/pre-final2-0M.txt  | awk '{print $6}' | rev | tr "[ATGC]" "[TACG]" | awk '{print "aaac"$0}'> Processing_files/pre-final2-0M.rev.txt
paste -d'\t' Processing_files/pre-final2-0M.txt Processing_files/pre-final2-0M.rev.txt | awk '{print $0"\t""ttgt"$6}' | awk 'BEGIN { OFS = "\t" } {gsub("aaacqeSediuC","Primer 1 (guide reverse)",$20) && gsub("ttgtGuideSeq","Primer 2 (guide forward)",$21)} 1' > $REFERENCE_GENOME"_LeishBASEv1_output.txt"

for i in $REFERENCE_GENOME"_LeishBASEv1_output.txt"
do
    mv "$i" "`echo $i | sed 's/.fasta//'`"
done

echo "LeishBASEedit v1 done"
```
